# Choline Transporter regulates olfactory habituation via a neuronal triad of excitatory, inhibitory and mushroom body neurons

**DOI:** 10.1101/2020.12.08.416891

**Authors:** Runa Hamid, Hitesh Sonaram Sant, Mrunal Nagaraj Kulkarni

**Affiliations:** Centre for Cellular and Molecular Biology, Council of Scientific and Industrial Research (CSIR-CCMB), Uppal Road, Hyderabad, India

**Keywords:** Mushroom body, *Drosophila*, Choline transporter, Habituation, Acetylcholine, GABAergic neurons, Cholinergic neurons, Autism spectrum disorder

## Abstract

Choline is an essential component of Acetylcholine (ACh) biosynthesis pathway which requires high-affinity Choline transporter (ChT) for its uptake into the presynaptic terminals of cholinergic neurons. Previously, we reported a predominant expression of ChT in memory processing and storing region of *Drosophila* brain called mushroom bodies (MB). It is unknown how ChT contributes to the functional principles of MB operation. Here, we demonstrate the role of ChT in non-associative form of learning, *Habituation*. Odour driven habituation traces are laid down in ChT dependent manner in antennal lobes (AL), projection neurons (PN) and MB. We observed that reduced habituation due to knockdown of ChT in MB causes hypersensitivity towards odour, suggesting that ChT also regulates incoming stimulus suppression. Importantly, we show for the first time that ChT is not unique to cholinergic neurons but is also required in inhibitory GABAergic neurons to drive habituation behaviour. Our results support a model in which ChT regulates both habituation and incoming stimuli through multiple circuit loci via an interplay between excitatory and inhibitory neurons. Strikingly, the lack of ChT in MB recapitulates major features of Autism spectrum disorders (ASD) including attenuated habituation, sensory hypersensitivity as well as defective GABAergic signalling. Our data establish the role of ChT in habituation learning and suggest that its dysfunction may contribute to neuropsychiatric disorders like ASD.

## Introduction

Acetylcholine (ACh) is the fundamental neurotransmitter of cholinergic neurons. These neurons are widely distributed throughout the central nervous system (CNS) in both vertebrate and invertebrate brain. In vertebrates, all pre-ganglionic sympathetic neurons, part of post-ganglionic sympathetic neurons, pre and post ganglionic parasympathetic neurons are cholinergic (Karczmar, 2007). Also, in invertebrates like *Drosophila,* almost all major types of sensory neurons including chemosensory, chordotonal, olfactory neurons and most regions of the central brain and interneurons are cholinergic (Grunewald and Siefert, 2019; Pereira et al., 2015; Salvaterra and Kitamoto, 2001). Given the widespread distribution of cholinergic neurons in the vertebrate and invertebrate brain, it is palpable that the ACh mediated neurotransmission is crucial for neural functions that include varied sensory modalities.

ACh synthesis for efficient neurotransmission at cholinergic synapses depends on the proteins of its metabolic cycle, namely, Choline acetyltransferase (ChAT), vesicular acetylcholine transferase (VAChT), Acetylcholine esterase (AChE) and ChT. Import of choline via ChT is the rate-limiting step of ACh biosynthesis. The MB in *Drosophila* brain are bilateral neuropilar areas which are principally cholinergic expressing ChAT and VAChT (Barnstedt et al., 2016; Croset et al., 2018). Based on immunostainings, we recently reported that there is a preponderance of ChT in *Drosophila* MB as compared to ChAT and VAChT (Hamid et al., 2019). The functional relevance of ChT expression in MB is not clear. MBs receive olfactory input from the antennal lobes (AL) via projection neurons (PNs) and serve as the prime site for sensory integration and learning. MB associate the memory with a reward or punishment pathway. However, before establishing such an association in MB, an animal must evaluate each incoming stimulus, identify the salient stimuli and initiate the appropriate stimulus driven response. Therefore, to understand the mechanisms of complex associative learning, it is important to decipher the mechanisms in MB that imparts an animal the flexibility to choose only the salient incoming stimuli, ignore inconsequential ones and finally register this information in a context dependent manner. Habituation learning enables an organism to evade such inconsequential stimuli. Several reported *Drosophila* genes like *dunce*, *rutabaga*, *turnip*, *radish*, *DCO*, *Leonardo*, *DAMB*, *Nmdar1 and 2* have high expression in MB and contribute to *Drosophila* memory (Keene and Waddell, 2007). Many of the proteins which are expressed at elevated levels in the MB are required for both associative learning as well as habituation (Duerr and Quinn, 1982; Engel and Wu, 2009) suggesting that the proteins contributing to associative learning also contributes to habituation. This implies the existence of an association for the two forms of plasticity. Flies that lack MB display reduced habituation (Acevedo et al., 2007; Cho et al., 2004; Duerr and Quinn, 1982; Roussou et al., 2019; Semelidou et al., 2018).

Habituation enhances attention only on salient features such as food, mate or danger in the surroundings. Habituation has been observed in many organisms from as simple as a single cell protozoa to *Aplysia californica*, medicinal leech, territorial fish, Birds and *Drosophila* to more complex life forms like rats, and humans suggesting its ubiquitous persistence (Eisenstein, 2001). Multiple habituation studies focused on *Drosophila* sensory system such as olfactory, gustatory, visual and proprioceptive system have contributed to our understanding of the cellular and circuit basis of habituation (Engel and Wu, 2009). At the synaptic level, habituation may result through the action of heterosynaptic modulation involving activation of inhibitory neurons or homosynaptic depression of excitatory neuron (Ramaswami, 2014). Studies in adult *Drosophila* suggest that potentiation of GABAergic inhibition onto PN terminals in AL cause olfactory habituation (Twick et al., 2014). Habituation reflects the efficient sensory input processing, and defective or premature habituation may lead to sensory hyper-responsivity which has been widely observed in ASD individuals (Foss-Feig et al., 2013; Gomot et al., 2002; Hudac et al., 2018; Kleinhans et al., 2009). Recently, orthologs of 98 human intellectual disability (ID) genes were reported to be important for habituation in fruitflies and a large fraction of these genes were associated with ASD (Fenckova et al., 2019). In view of our previous findings that report high expression of ChT in MB, we study a putative role of ChT in ‘Habituation’ which is widely regarded as a prerequisite for more complex form of associative learning. Compilation of the previous information describing habituation in *Drosophila* reveals that most of the olfactory habituation paradigms engage olfactory circuitry as well as mushroom body intrinsic neurons (Das et al., 2011; Larkin et al., 2010; Paranjpe et al., 2012; Roussou et al., 2019). Therefore, we mapped the function of ChT in MB neurons as well as in the olfactory pathway governing olfactory habituation learning in *Drosophila* larvae.

Here, we show that ChT is required for habituation plasticity at multiple loci involving olfactory pathway and MB neurons. This study report for the first time that ChT is also localised in GABAergic terminals present in MB calyx, suggesting that ChT is not unique to cholinergic neurons. We show that both excitatory and GABAergic inhibitory synapses co-recruit ChT in the MB calyx and their teamwork regulate habituation as well as its central operational features: the response devaluation to an olfactory stimulus, spontaneous recovery on the removal of stimulus or dishabituation of response upon exposure to an unrelated stimulus. Knock-down of ChT in MB was observed to be correlated with the hypersensitivity towards the incoming stimuli suggesting ChT bridges the link between upstream plasticity and downstream stimulus suppression for habituation. Our results demonstrate the role of a conserved protein, ChT, contributing to the dynamic nature of habituation learning and highlight that its dysfunction leads to sensory abnormalities similar to observed in ASD.

## Material and methods

### *Drosophila* stocks and maintenance

All *Drosophila* stocks were grown on standard BDSC formulation of fly food consisting of cornmeal-agar media supplemented with yeast, grown at 25°C unless mentioned otherwise on a 12:12 light:dark cycle. For RNAi experiments, all fly crosses and their genetic controls were grown at 29°C. For control experiments, all driver, as well as responder lines were crossed with *w^1118^* (+). For temperature-sensitive experiments requiring GAL80, the crosses were maintained at 18°C and shifted to 29°C before the experiments. The Gal4 drivers *MB247GAL4* (#50742), *c305aGAL4* (#30829), *Or83bGAL4* (#26818), *GH298GAL4* (#37294), *GH146GAL4/CyoGFP* (#30026), *GAD1GAL4* (#51630), *ChATGAL4* (#6798), *TubGAL80^ts^* (#7017) were obtained from Bloomington stock centre, Bloomington, Indiana. The UASRNAi strains for ChT (101485) was obtained from VDRC and BDSC (#28613). NP1131 (103898) was from DGRC Kyoto stock centre, Japan. UAS-ChT line was previously created and characterised (Hamid et al., 2019). All genetic combinations of stocks used in this study were created in the lab.

### Antibodies and immunohistochemistry of larval brain

For immunostaining experiments, the larval brains were dissected in phosphate buffer saline (PBS), fixed in 4% paraformaldehyde followed by 5 washes with PBS+ 0.5% TritonX-100 (PBSTx) for 6 min each. The brains were incubated with 5% BSA for 40 mins and incubated at 4°C with primary antibody for 16 hrs. The primary antibody was removed followed by 5 washes for 5 min each with PBSTx at room temperature. The brains were then incubated with secondary antibody for 1 hour followed by 3 washes with PBSTx of 10 min each. The immunostained samples were mounted in vectashield and confocal images were collected using Zeiss LSM880 microscope. Rabbit *Drosophila* anti-ChT was used at 1:300 and mouse anti-DLG (DSHB) was used at 1:200. Optical slices were obtained in 1024×1024 pixel format using oil immersion 40 x (1.35 N.A) or 63x (1.4 N.A) objectives, Imaging facility, CCMB, India. The represented images were stack projected using maximum intensity projection module in Image J 1.52p (NIH, USA).

### Measurement and quantification of olfactory habituation index

The assays were performed using 90 mm Petri dishes with 2% solid agar medium. The plates were divided into two halves and two 1.5ml eppendorf caps were cut, autoclaved and embedded into the agar on opposite sides of the plates to place the odour. Only third instar foraging larvae (approx. 72 – 90 hr AEL) were used for the experiments, as they are continuously feeding and their tendency to move towards the fruity odour is higher. The assays were performed at room temperature and under red light to avoid any visual inputs. Also, all experiments were performed before noon to avoid any alterations arising due to different activity phases of larvae during the day. Thirty to forty foraging larvae were thoroughly washed and placed in the centre of the agar plate. Immediately, the odour was placed near one end of the plate and water at the opposite end. After three minutes the number of larvae on each half of the agar plate was counted. The quantification used the midline drawn in the centre of the plate to allocate whether the larvae were on the odour side or the water side. The response towards the odour was measured by calculating the response index (R.I) and mentioned as pre-R.I:

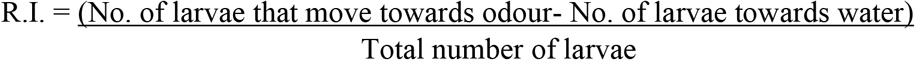

The larvae were allowed to crawl on the plate in presence of odour for a total of five to six minutes to allow them to habituate with the odour. They were then placed back in the middle of the agar plate and response index was calculated again and mentioned as Post R.I.

For quantifying habituation, the Habituation index (H.I) was calculated as:

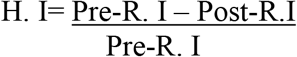

We used pure ethyl acetate (ETA), 1:100 dilution of Amyl acetate (AMA) and 1:1000 dilution of 3-Octanol.

### Analysis of spontaneous recovery and dishabituation

Thirty to forty larvae were habituated in presence of the odour for five to six minutes on an agar plate. Subsequently, the habituated larvae were immediately placed on a separate agar plate in the absence of odour for recovery. The response towards the odour was measured after 15, 30 and, if required, 60 min. R.I was calculated as mentioned above and referred to as recovery-R.I. The calculated response index after 15 or 30 min (recovery-R.I) was compared with the post-R.I to assess for the spontaneous recovery.

For Dishabituation, the larvae were habituated for five to six minutes and were subsequently given a cold shock for 1 min by placing them on a pre-cooled agar plate on ice for one minute. The larvae were then kept for 2 min on agar plate without odour at room temperature to regain their movement caused due to low temperature followed by response index test. R.I towards odour was calculated as mentioned earlier and was compared with the post-R.I for the odour to assess for dishabituation.

### Statistical Analysis

Statistical analysis was performed using GraphPad Prism version 8.0.0 for Windows, GraphPad Software, San Diego. Each data set was compared with their genetic controls which is indicated separately in individual figure legend. The significance of difference was assessed using Mann-Whitney U test and computing exact p-values. Data was considered significantly different if p<0.05 and represented as *** for p<0.0001, ** for p<0.001, * for p< 0.01 and n.s for non-significance when p>0.05

#### Model

Larval anatomical schematics and Figure 7 were prepared using Biorender (www.biorender.com)

## Results

### Decrement of odour-specific chemotactic response in naïve *Drosophila* larvae conforms to habituation parameters

Attraction towards an odour is referred to as chemotaxis which is essential for a diversity of insects to navigate for food sources, potential mating partner, assessing danger, search for egg-laying sites. The olfactory system of insects has evolved to impart them great discriminatory power for behaviorally relevant odours, decipher this message in CNS and finally exhibit appropriate behaviours. Thus, chemotaxis based behavioural responses towards an odour are immensely important for their survival. Foraging *Drosophila* larvae are in constant search of food. The larval olfactory system is similar but numerically simpler than the adults, discern a wide range of odours and can learn to discriminate between different odours as well as concentration (Khurana and Siddiqi, 2013). Thus, it provides us with a system that has a genetically accessible well-defined neural circuit relevant for our study.

We first tested and standardised the assay in wildtype 3^rd^ instar foraging larvae to affirm if our assay accedes to habituation parameters. Continuous exposure of naïve wild type larvae for 5 min to ETA and AMA, evoked significant avoidance of these odours (Fig. 1 A). ETA elicited the strongest chemotactic response as well as showed the strongest avoidance after prior exposure among both odours tested. Therefore, we used the attractant ETA for subsequent experiments. We further investigated whether the decrement of chemotactic response conforms to classical habituation parameters (Rankin et al., 2009). This means that animals that habituate to a stimulus should regain the response after a time-lapse, if the stimulus is withheld. This phenomenon is termed as ‘spontaneous recovery’. Indeed, we observed a spontaneous recovery of the chemotactic response to the naïve levels after 15 and 30 min rest time (Fig. 1A). To confirm the initial decrement as habituation and not a fatigue or sensory adaptation, we tested another classical feature of habituation, called dishabituation.

**Figure 1:**
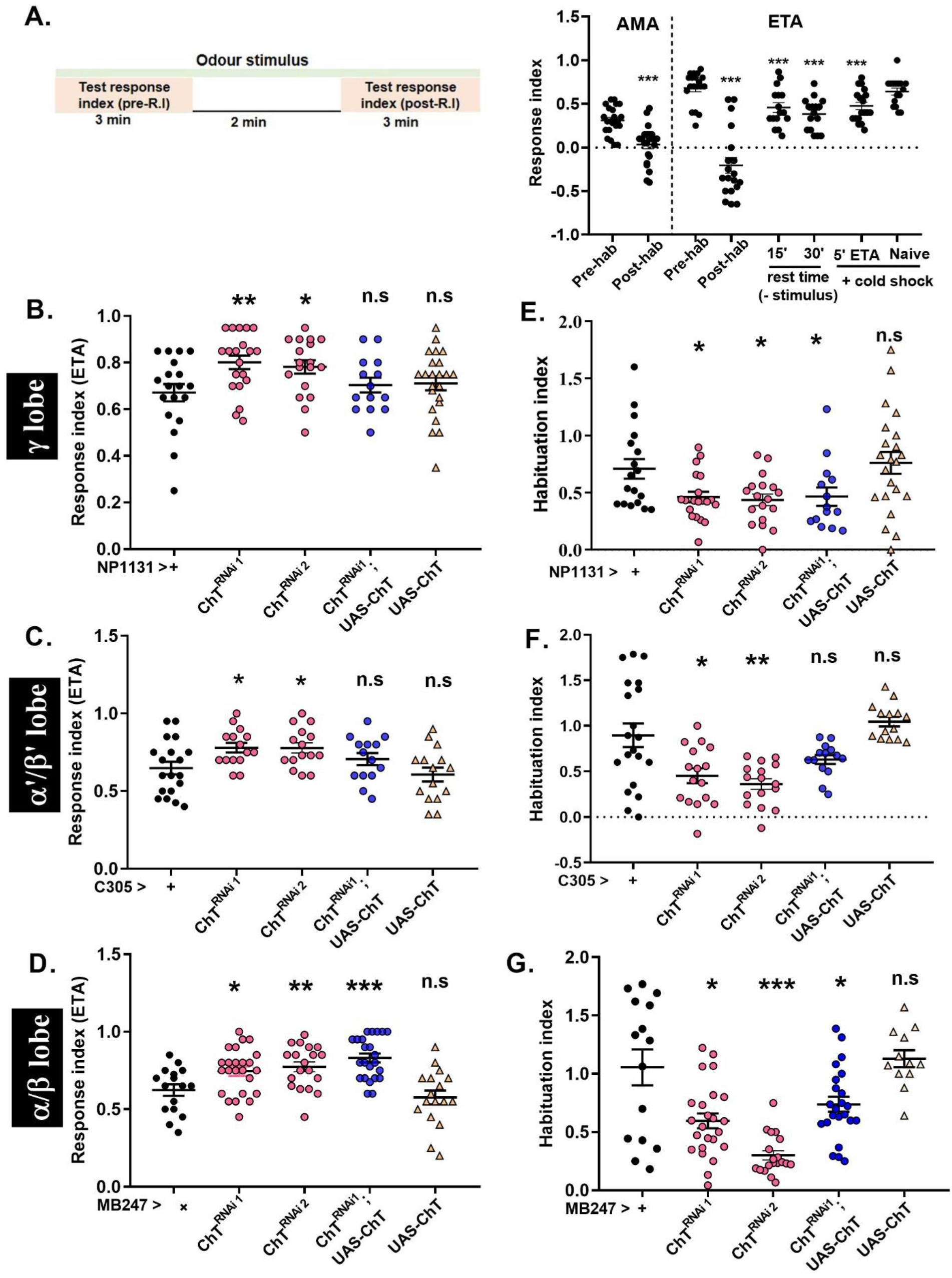
Knock-down of ChT in MB intrinsic neurons enhances chemotaxis towards odour but supresses habituation. (A) Schematics represent the specific time segments followed for the experiments. Scatter plot shows response indices (R.I, black circles) of naïve wild type larvae (W^1118^) towards amyl acetate (AMA) or ethyl acetate (ETA) (pre-hab), after 5 min of odour exposure (post-hab). R.I, following a 5 min exposure to ETA, after 15 or 30 min in absence of stimulus (recovery) and after 1 min cold shock (Dishabituation, 5’ETA+cold shock). Naïve+cold shock represents the data of cold shock given to larvae without odour pre-exposure; N≥16. *(B-D)* response index of naïve larvae, and *(E-G)* habituation index of larvae exposed to ETA of genotypes: *NP1131*> *ChT^RNAi1^* or *ChT^RNAi2^ and NP1131> ChT^RNAi1^;UAS-ChT, NP1131*> *UAS-ChT* as compared to their controls *NP1131*> *+* (B and E), *C305a*> *ChT^RNAi1^* and *ChT^RNAi2^*, *C305a*> *ChT^RNAi1^;UAS-ChT, C305a*> *UAS-ChT* as compared to their controls *C305a*> *+* (C and F), *MB247*> *ChT^RNAi1^* and *ChT^RNAi2^*, *MB247*> *ChT^RNAi1^;UAS-ChT, MB247*> *UAS-ChT* as compared to their controls *MB247*> *+*(D and G). Pink circles represent knock down using *UAS-ChT^RNAi 1^* and *UAS-ChT^RNAi2^*, Blue circle represent rescue, and yellow triangles represent transgenic over-expression of *UAS-ChT* as compared to genetic controls (black circles). Data is represented as scatter plot with error bars showing SEM and N≥16. Each N represent one experiment performed with group of 30-40 larvae. Statistical significance was determined by Mann-Whitney U test and *** represent p≤0.0001, n.s means statistical non-significance with p≥0.05. Also, refer to supplementary figure S5 for comparisons of R.I and H.I of heterozygous combinations of *MBGAL4s* >+ versus *UAS-ChT^RNAi 1^* >+ and *UAS-ChT^RNAi2^* >+

It means if the habituated animal is exposed to an unrelated strong stimulus, the naïve response is fully/ partially restored. We attempted dishabituation using a 1 min exposure to cold shock on ice. This significantly reverses the chemotactic response of larvae pre-exposed to 5’ ETA (Fig. 1A). Importantly, 1 min exposure of cold shock given to naïve larvae had no effect on chemotaxis towards ETA suggesting cold shock does not affect the general perception of olfactory stimuli but causes loss of habituation (dishabituation), Fig. 1A. The decrease in response towards the ETA after pre-exposure as well as the naïve response cannot be attributed to the locomotor defect because most of the larvae left the choice point.

Taken together, the decrement of chemotaxis in wildtype larvae on continuous exposure to an odorant stimulus, the recovery of the response when the stimulus was withheld as well as the dishabituation conforms to the habituation parameters and demonstrate that the response attenuation is attributed to olfactory habituation. We have used these olfaction-based paradigms in our subsequent experiments to demonstrate the functional relevance of ChT in regulating habituation learning.

### Intrinsic neurons of MB require ChT for olfactory habituation learning and odour stimulus suppression

There are three types of MB intrinsic neurons i.e, α/β, α’/β’ and ϒ neurons, also termed as Kenyon cells (KC). These are distinctly implicated in olfactory learning, memory (Chia and Scott, 2020; Krashes et al., 2007). To investigate the functional significance of ChT in MB we depleted it in independent domains of MB intrinsic neurons with the help of *UAS-GAL4* binary expression system (Brand and Perrimon, 1993) and asked if these neurons require ChT function in habituation learning. For this, two RNAi fly stocks [ChT^RNAi1^ (V101485) and ChT^RNAi2^ (BL28613)] were used to knock-down ChT in MB intrinsic neurons. The knockdown efficiency of ChT^RNAi2^ was assessed by immunostaining (Fig. S1) while of ChT^RNAi1^ was described previously by us (Hamid et al., 2019). First, we assessed the naïve chemotactic response of 3^rd^ instar foraging larvae towards ETA upon knockdown of *ChT* in each of the neuronal domain of MB. Interestingly, we observed a significant increase in naïve chemotactic response upon knockdown of ChT in ϒ lobe neurons (NP1131> *ChT^RNAi1^ or ChT^RNAi2^*) as well as in α’/β’ (C305a> *ChT^RNAi1^ or ChT^RNAi2^*) and α/β (MB247> *ChT^RNAi1^ or ChT^RNAi2^*) class of KC as compared to their genetic controls (MBGAL4s > +; Fig 1B-D). To test if this enhancement is specific to the knockdown of ChT, we expressed ChT transgene on *UAS-ChT^RNAi1^* background in all the three neuronal domains of MB (*MBGAL4s> ChT^RNAi1^;UAS-ChT*) and observed a reversal of the response index in ϒ and α’/β’ class of KC but not in α/β class of KC (Fig 1B-D). Next, we tested the effect of ChT knockdown in MB on habituation learning. A significant reduction in habituation index was observed in the group of larvae upon reduction of ChT with both *ChT^RNAi1^ and ChT^RNAi2^* fly lines in ϒ, α’/β’ and α/β KC (Fig.2 E-G). Transgenic expression of ChT in *UAS-ChT^RNAi1^* background significantly enhanced the habituation index of the larvae in α’/β’ but not in ϒ and α/β KC (Fig.2 E-G). This difference in the rescue of response index as well as habituation index in different KC may be due to the differences in the expression levels of *UAS-ChT^RNAi1^* and *UAS-ChT* transgenes driven by the different GAL4s. To clarify, if the levels of ChT determine the extent of habituation and chemotaxis, we over-expressed ChT in all the three classes of KC neurons using specific GAL4 drivers (MBGAL4’s>*UAS-ChT*). The extent of the naïve olfactory response, as well as the habituation, remains unaffected by overexpression of ChT (Fig.2 B-G). Next, we confirmed if the hypersensitivity to ETA and decreased habituation is specific to the knockdown of ChT and not specific to the kind of pre-exposed odour. We knocked down ChT in ϒ, α’/β’ and α/β KC using *UAS-ChT^RNAi^*^1^ (*MBGAl4s>ChT^RNAi1^*) and exposed these group of larvae to amyl acetate (AMA) and the alcoholic class of odour, 3-Octanol. For both the pre-exposed odours, we observed an enhancement of chemotaxis and reduced habituation suggesting it to be ChT specific phenotype and not to the class of odour (Fig. S2 A-D). To further ascertain that attenuation of naïve chemotactic response and habituation index is indeed due to knockdown of ChT and not due to non-specific secondary effect of the transgenic lines, we also examined the heterozygous combination of *UAS-ChT^RNAi1^* and *UAS-ChT^RNAi2^* with wildtype lines (*UAS-ChT^RNAi1^* >+ or *UAS-ChT^RNAi2^* >+) and did not observe any significantly different response compared to the different MBGAL4s>+. (Fig. S5).

**Figure 2:**
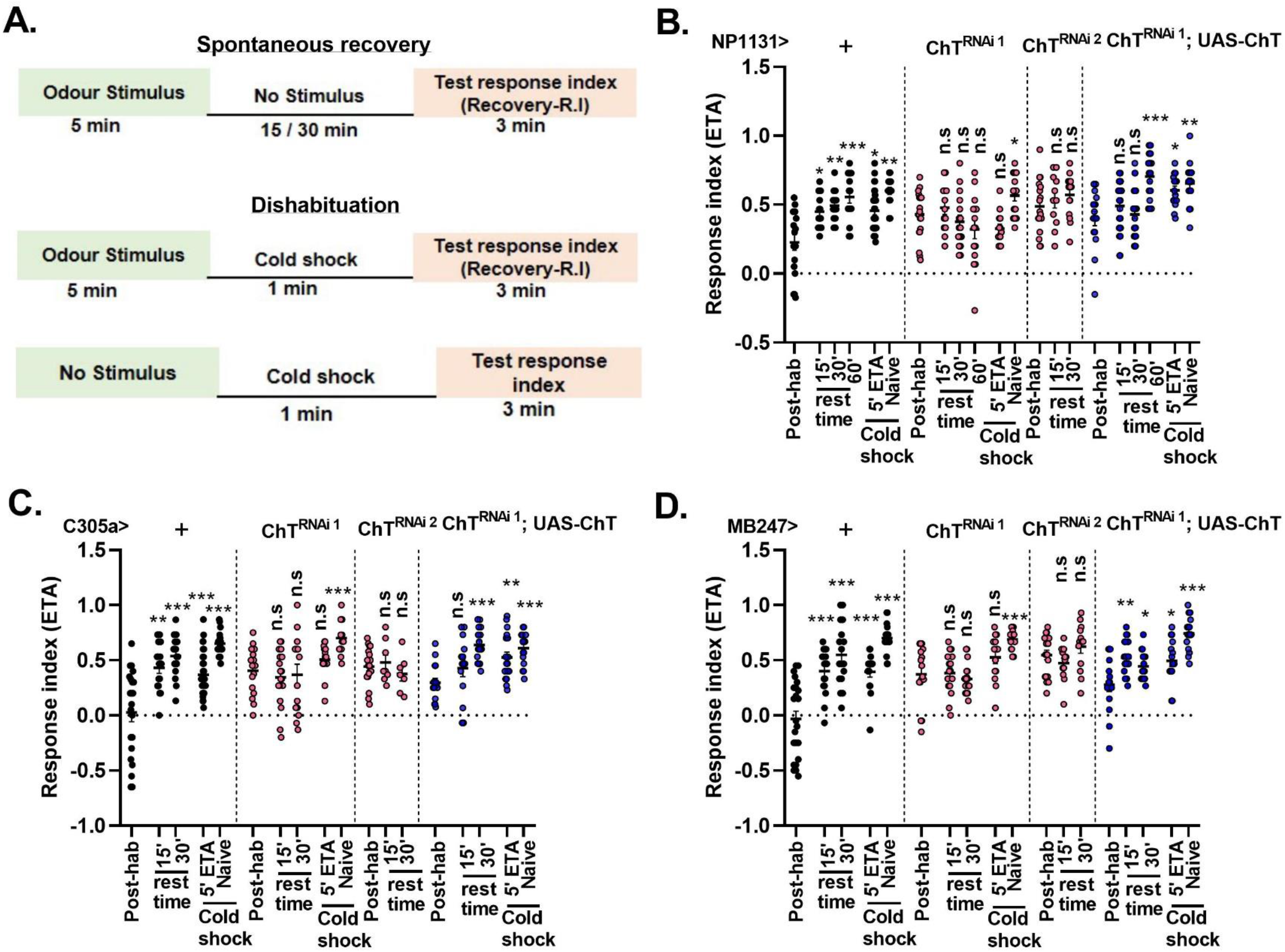
Knockdown of ChT in MB intrinsic neurons perturb spontaneous recovery and dishabituation of habituated larvae. (A) Schematics of time segment followed for assessing spontaneous recovery and dishabituation. (B-D) Scatter plot shows response indices towards ETA of larvae exposed to 5 min ETA (Post-hab), in absence of stimulus after 15, 30 or 60 min (rest time) presented to 1 min cold shock (5’ETA+coldshock), naïve larvae presented to 1 min cold shock (naïve + cold shock) of genotypes: (B) control *NP1131*> *+ (*ϒ lobe; black circles, N≥16), *NP1131GAL4> ChT^RNAi1^ or ChT^RNAi2^* (pink circles, N≥11), rescue *NP1131GAL4> ChT^RNAi1^;UAS-ChT* (blue circles, N≥16) (C) control *C305aGAL4*> *+ (*α’/β’lobe, black circles, N≥16), *C305aGAL4> ChT^RNAi1^ or ChT^RNAi2^* (pink circles, N≥9), rescue *C305aGAL4> ChT^RNAi1^;UAS-ChT* (blue circles, N≥14). (D) control *MB247GAL4*> *+ (*α/β lobe, black circles, N≥17), *MB247GAL4> ChT^RNAi1^ or ChT^RNAi2^* (pink circles, N≥13), rescue *MB247GAL4> ChT^RNAi1^; UAS-ChT* (blue circles, N≥16). Statistical significance was determined by Mann-Whitney U test and *** represent p≤0.0001, n.s means statistical non-significance with p≥0.05.

Previously, we reported attenuated neuromuscular junctions (NMJ) in third instar larvae caused due to knockdown of ChT in α/β and ϒ intrinsic lobes of MB (Hamid et al., 2019). To ascertain if the observed reduction in habituation index is the acute functional change in the neurons during habituation or have resulted due to changes in the NMJ, we employed the TARGET system (McGuire et al., 2004). TARGET uses a Tubulin promotor to express temperature-sensitive repressor of GAL4 (TubGAL80^ts^) where expression of GAL80^ts^ allows RNAi expression by GAL4 at 29°C but not at 18°C. Using this system, we limited the expression of *UAS-ChT^RNAi1^*(*MBGAL4>ChT^RNAi1^;tubGAL80^ts^*) to the developmental window of foraging 3^rd^ instar larvae during which the assay was performed. We observed a significant enhancement in response index towards the odour (Fig. S3A) and a drastic reduction in habituation index (Fig. S3B) upon knockdown of ChT in α/β and ϒ lobe neurons at 29°C but not at 18°C (Fig. S3 A and B). These results confirm that ChT is acutely required in MB intrinsic neurons to facilitate habituation and the observed habituation defects are not due to changes in neurons during development. Altogether, our results indicate that knockdown of ChT in MB does not affect the odour perception. It has a function in MB intrinsic neurons that support devaluation of the incoming stimuli and regulates sensitivity towards incoming olfactory stimuli.

### ChT is essential in intrinsic neurons of MB for the maintenance of key characteristics of habituation

A habituated animal should reestablish the response towards stimuli partially or wholly within a specific time frame if the stimulus is withheld (Rankin et al., 2009). To ascertain that the decrement in response due to knock down in ChT is indeed habituation phenotype, we assessed the response recovery (spontaneous recovery), after a rest period of 15 min and 30 min, in the habituated larvae when the stimulus was withheld. The knockdown of ChT using the expression of *UAS-ChT^RNAi1^* and *UAS-ChT^RNAi2^* lines driven by *NP1131GAL4* (ϒ lobe), *C305aGAL4* (α’/β’lobe) and *MB247GAL4* (α/β lobe) intrinsic lobes caused a defect in the larvae to spontaneously recover (Fig. 2 B-D). On the other hand, larvae from their genetic controls (*MBGAL4s*>+) significantly recovered to the naïve levels just after 15 min (Fig. 2 B-D). The defect in recovery due to depletion of ChT recovered partially or completely when *ChT* was transgenically expressed on a *ChT^RNAi1^* background in α’/β’, α/β and ϒ class of KC neurons (Fig. 2 B-D). However, we noticed a complete recovery in ϒ lobe only after 60 min whereas in α’/β’ and α/β the recovery was achieved within 30 min.

Next, we induced dishabituation by exposing the habituated larvae to cold shock for 1 min. The knockdown of ChT in the ϒ, α’/β’ and α/β class of KC neurons of larvae using *NP1131GAL4* (Fig. 2B)*, C305aGAL4* (Fig. 2C) and *MB247GAL4* (Fig. 2D) driver lines were unable to dishabituate upon cold shock. On the other hand, exposure of habituated larvae to 1 min cold shock reverses the decrement of chemotaxis to naïve levels in the control group of larvae (*MBsGAL4*>+; Fig. 2 B-D). The dishabituating capability was rescued back in these larvae when ChT was transgenically expressed on a ChT^RNAi^ background in ϒ, α’/β’, and α/β class of KC neurons (Fig. 2 B-D). The exposure of naïve larvae to similar cold shock did not affect the chemotaxis, suggesting the observed enhancement of response index is not the sensitisation of olfactory receptors due to cold shock.

The control group of larvae (*MBsGAL4*>+) are capable of reestablishing baseline response levels that follow an unrelated dishabituating stimulus as well as after a prolonged lapse of time in absence of stimulus while larvae deficit of ChT function in MB neurons are incapable of resuming the response levels. This suggests that in absence of ChT, the synaptic capability to re-establish the response is compromised leading to defective spontaneous recovery and dishabituation. These results strongly suggest that ChT potentially contributes to synaptic plasticity that results in olfactory habituation learning through MB intrinsic neurons.

### ChT is localized in the neural circuit of the larval olfactory pathway

*Drosophila* larva has a dome of perforated cuticle called dorsal organ (DO) at its anterior end through which odours can pass (*schematics* Fig. 3A). There are two bilaterally symmetrical DO in *Drosophila* larva each housing 21 olfactory sensory neurons (OSN) (Python and Stocker, 2002). The axons of OSNs bundle together and project into the AL via an antennal nerve. Each OSN expresses a specific type of olfactory receptor (OR) along with the OR83b gene which is essential for the proper functioning of ORs (Larsson et al., 2004). OSN responds to the odours by the combinatorial activity of several OSN types. The terminals of OSNs ramify into branches of different density and size (Masuda-Nakagawa et al., 2009). We investigated the presence of ChT at OSN terminals and visualised them using *Or83bGAL4* to drive *UAS-mCD8GFP* and costained these terminals by the *anti-* ChT antibody. We observed ChT expression at few of the termini of its branches. It was not as profusely defined as the total number of branches suggesting that its function lies in a particular subset of OSNs (Figure S4. A). OSNs project onto the local interneurons in AL or dendrites of projection neurons in the antennal lobes (*schematics* Fig. 3A). Projection neurons (PN) are uniglomerular and relay information to mushroom bodies and other higher brain regions (Tanaka et al., 2012). They connect to individual glomeruli of calyx via the presynaptic ends and the AL via postsynaptic ends (Masuda-Nakagawa et al., 2009). We used *GH146GAL4* to drive expression of *UASmCD8GFP* reporter in projection neurons. The dendritic arborization of PN covered a larger area of the antennal lobe (Fig. 3B). An axon extending from AL was visible and terminate in the MB calyx which represents the output area of PN (Fig 3B.). We observed an intense localisation of ChT at the presynaptic ends of PN terminating in the calyx of MB (Fig 3C.). The dendritic end showed a sparse localisation of ChT which probably represent the expression of ChT in the antennal lobes rather than the dendritic end of PNs (Fig. 3D). To identify ChT localisation in the larval antennal lobe neurons, we crossed *GH298GAL4* driver with *UAS-mCD8GFP* and coimmunostained with anti-ChT to visualize ChT expression pattern in AL neurons (Fig. 3 E-G). The *GH298*GAL4 expression was seen as arborisation in the antennal lobes and number of cells situated around the lateral edge of the AL (Fig. 3E). GH298 expression was also seen in MB calyx with a predominant colocalization with anti-ChT immunostainings (Fig. 3H-J). Collectively, our immunostaining data shows that ChT is present in all three major cellular components of the olfactory pathway, the OSNs, the AL and the MB of *Drosophila* larva. Therefore, it became logical to investigate the role of ChT in the whole olfactory neural circuitry and assess if ChT contributes at each stage of olfactory processing to facilitate olfactory habituation.

**Figure 3:**
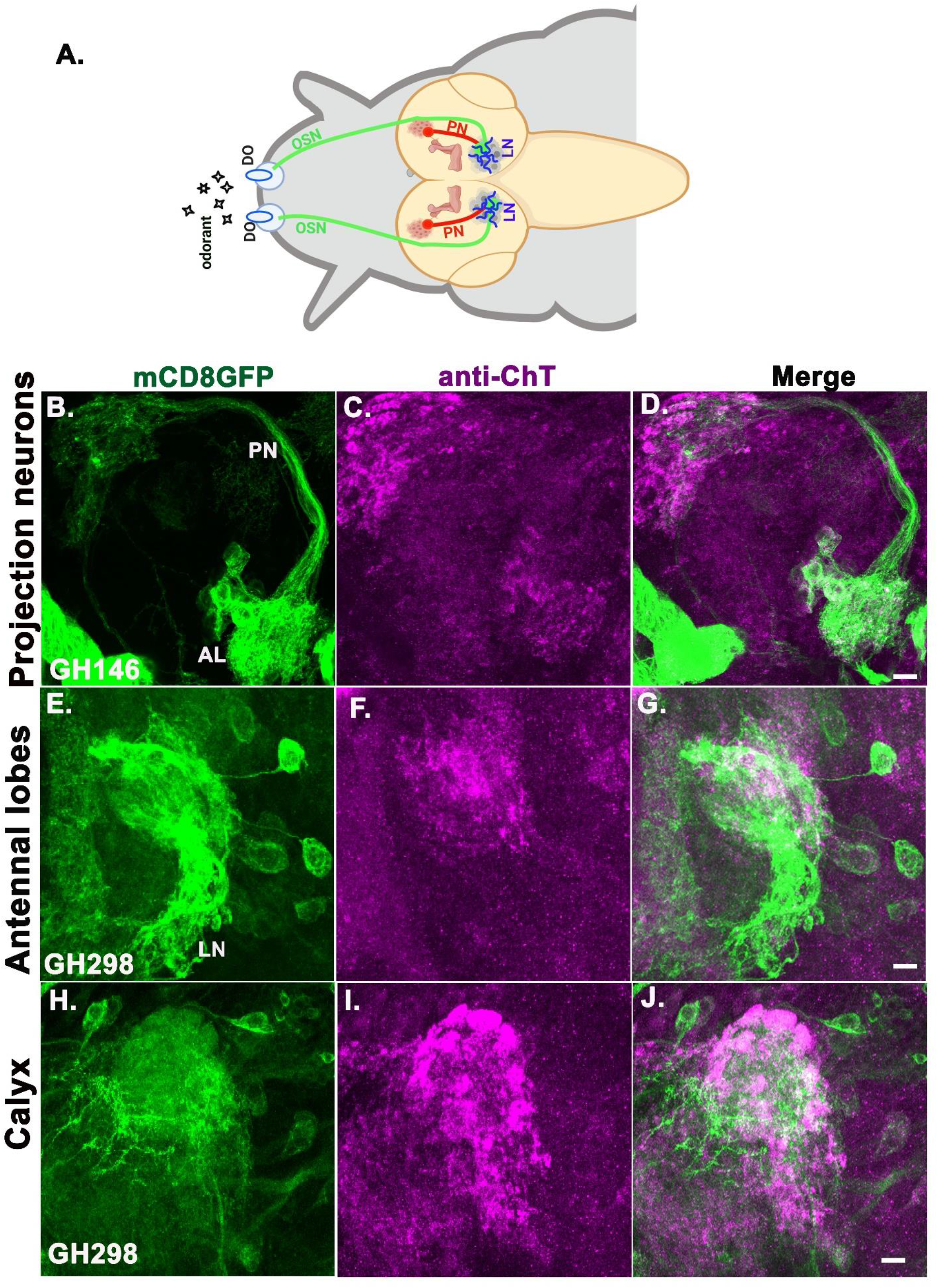
Localization of ChT in the larval olfactory circuit: (A) Schematics of larval olfactory pathway. Odorants are detected by OSN (green) which terminate in AL and synapses with LN (blue) or with dendrites of PN (red). PN innervate MB calyx (brown). (B-D) Expression of *UAS-mCD8GFP* in projection neurons (green) driven by *GH146GAL4* and immunostained with anti-ChT (magenta) (E-G) antennal lobes, and (H-J) calyx driven by *GH298GAL4>mCD8GFP* and immunostained with anti-ChT (magenta). Colocalized regions of mCD8GFP and ChT in panel D, G and J are merged as white. All images are z-stacked pseudocoloured representative of 3-5 larval brains. Scale bar 50 μm

### *ChT* mediates habituation learning but not chemotaxis through local interneurons of antennal lobes and projection neurons

Our finding that ChT is localised in olfactory circuit neurons, provided the basis to explore if functioning of ChT in olfactory circuit may potentially contribute to habituation learning. Therefore, to investigate the neural circuitry through which ChT could contribute to habituation learning, we first knocked down ChT in OSN by expressing *UAS-ChT^RNAi^*^1^ using *Or83b*GAL4 (*OR83b*GAL4>UAS-*ChT^RNAi^*) and observed a reduction in chemotaxis towards ETA as compared to their genetic control (*OR83b*GAL4>+) (Fig.S4. B). The attenuated chemotaxis was partially rescued by transgenically expressing *UAS-ChT* on a *ChT^RNAi^* background (*OR83bGAL4>UAS-ChT^RNAi^; UAS-ChT*) (Fig.S4. B). However, we did not find any significant reduction in habituation index upon knockdown of ChT in OSNs (Fig.S4.C). The overexpression of ChT in OSNs neither modulated chemotaxis nor the habituating capability of larvae (Fig. S4 B, C). These results imply that knockdown of ChT in OSNs impair only the chemotaxis.

When the larvae sense an odour through OSNs, an activity pattern is generated at the glomerulus of AL because the axons of OSNs either synapse with local interneurons (LN) in AL or with dendrites of the second-order neurons called PNs terminating in the AL. AL are referred to as the first olfactory processing centre in the brain. Our immunostaining analysis shows an extensive localisation of ChT in AL (Fig 3E-G). Therefore, we tested naïve chemotactic response as well as habituation index after knockdown of ChT in AL neurons using *GH298GAL4* driver line. We observed that depletion of ChT by *UAS-ChT^RNAi1^* and *UAS-ChT^RNAi2^* in AL neurons (*GH298> UAS-ChT^RNAi1^ or UAS-ChT^RNAi2^)* significantly reduced habituation as compared to their genetic controls (*GH298*> +) without affecting naïve chemotactic responses (Fig. 4A and B). The control larvae recovered back to the naïve levels within 30 min after stimulus removal (Fig. 4C). However, we observed that the response towards odour recovered to near baseline level when ChT was knocked down with *UAS-ChT^RNAi1^* but not with *UAS-ChT^RNAi2^* (Fig. 4C). The observed recovery obtained with *UAS-ChT^RNAi1^* might be because the decrement in chemotaxis was already less upon habituation. Also, the brief dishabituating cold shock stimulus recovers the response index to the naïve levels significantly in control (*GH298*> +) but not when ChT was knocked down in *GH298*>*ChT^RNAi1^* larvae (Fig. 4C). The failure of group of larvae with depleted ChT to spontaneously recover and dishabituate was significantly rescued by transgenic expression of ChT on ChT^RNAi1^ background (*GH298*> *ChT^RNAi1^;UASChT*) (Fig. 4 C). Since LNs modulate the activity of PNs and our immunostainings show ChT expression in PN presynaptic terminals, we assessed the functional role of ChT in PNs in odour processing and habituation. For this, we used *GH146GAL4* driver to drive *UAS-ChT^RNAi1^* and *UAS-ChT^RNAi2^* for knockdown of ChT (*GH146> UAS-ChT^RNAi1^ or UAS-ChT^RNAi2^)*. Knockdown of ChT in PNs did not cause any significant changes in naïve chemotactic response (Fig. 4D) but caused a significant reduction in habituation index with both ChT^RNAi1^ and ChT^RNAi2^ lines as compared to the genetic controls (*GH146GAL4*>+; Fig. 4E). The habituation defect was rescued back by transgenically expressing *UAS-ChT* in larvae of genotype *GH146*> *ChT^RNAi1^;UAS-ChT* (Fig. 4E). The decline in the olfactory response was recovered back to the baseline levels in the control group of larvae (*GH146*>+) over time, in absence of a stimulus (Fig. 4F) as well as on providing dishabituating cold stimulus conforming to the habituation characteristics (Fig. 4F). This spontaneous recovery and dishabituation were not observed in the group of larvae with depleted ChT in genotypes *GH146*>*ChT^RNAi1^ or ChT^RNAi2^* (Fig. 4 F).

**Figure 4:**
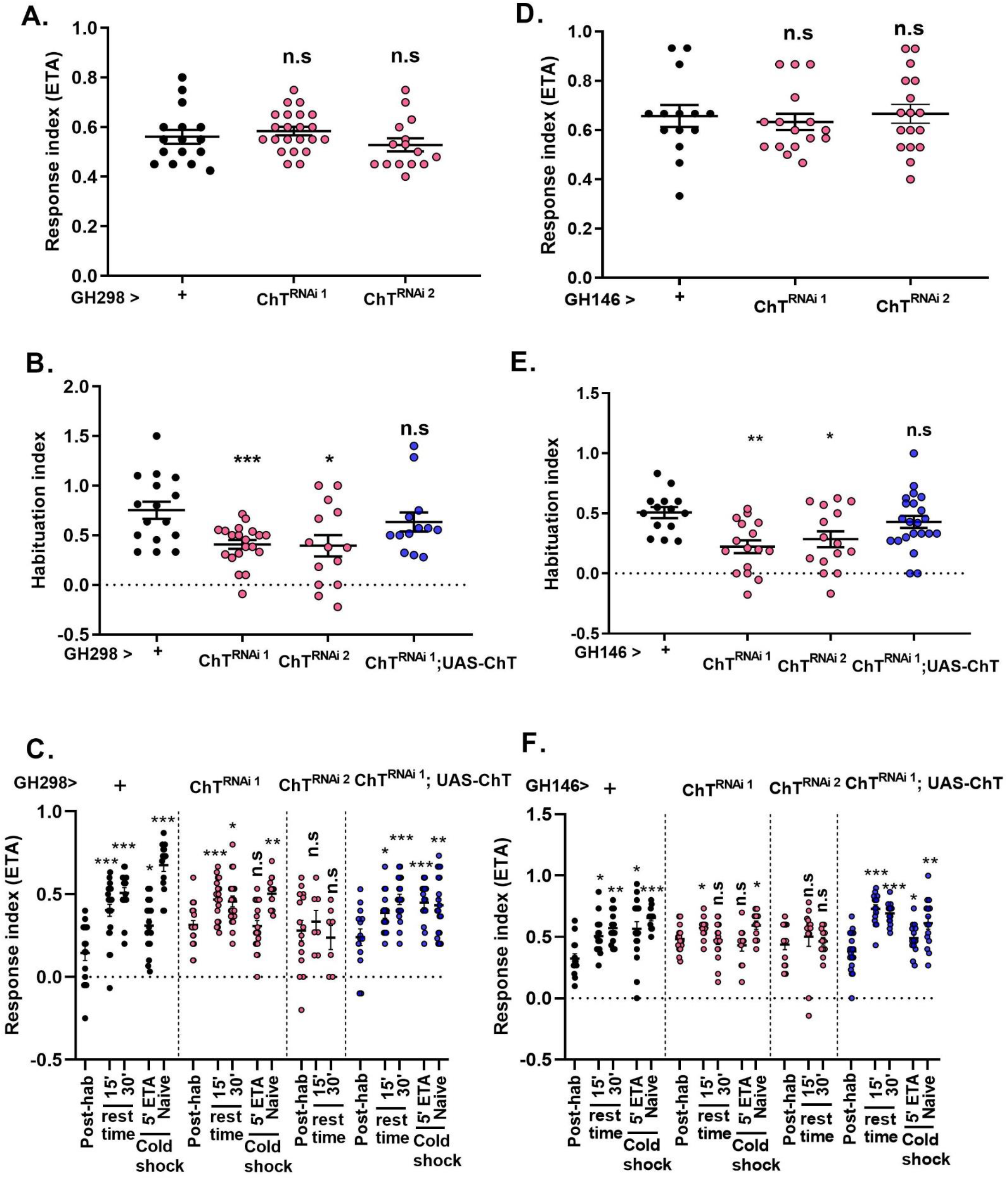
Presence of ChT in antennal lobes and projection neurons is required for habituation but not for chemotaxis. (A) Response index and (B) habituation index (C) Response index following 5’ ETA exposure (post-hab), spontaneous recovery after 15 and 30 min rest time in absence of stimulus, post-hab larvae presented to 1 min cold shock (5’ETA+coldshock) and naïve larvae presented to 1 min cold shock (naïve + cold shock), of genotypes: *GH298GAL4*> *+* (black circles, N≥16), *GH298GAL4>ChT^RNAi1^ or ChT^RNAi2^* (pink circles, N≥18), *GH298GAL4*> *ChT^RNAi1^;UAS-ChT (Blue circles,* N≥15). (D) Response index and (E) habituation index (F) response index following 5’ ETA exposure (post-hab), spontaneous recovery after 15 and 30 min rest time in absence of stimulus, post-hab larvae presented to 1 min cold shock (5’ETA+coldshock) and naïve larvae presented to 1 min cold shock (naïve + cold shock), of genotypes: *GH146GAL4*> *+* (black circles, N≥12), *GH146GAL4>ChT^RNAi1^ or ChT^RNAi2^* (pink circles, N≥12), *GH146GAL4*> *ChT^RNAi1^;UAS-ChT (Blue circles,* N≥14). Data is represented as scatter plot with error bars showing SEM and each dot represent one experiment performed with group of 30-40 larvae. Statistical significance was determined by Mann-Whitney U test and *** represent p≤0.0001, n.s means statistical non-significance with p≥0.05. Also, refer to supplementary figure S5.

Altogether, our results suggest that ChT plays a role in distributed plasticity including AL neurons, PNs and MB intrinsic neurons to facilitate olfactory habituation. Our results highlight the dynamic engagement of ChT in first processing and higher processing centres of the larval brain to regulate responses towards stimuli that are inconsequential in nature and function in a cooperative manner to optimise synchronisation of odour perception, information transfer and display relevant behavioural output.

### ChT is required in GABAergic neurons for both olfactory habituation and stimulus suppression but mediates only stimulus suppression via cholinergic neurons

OSNs and major class of PNs in *Drosophila* are cholinergic in nature (Yasuyama and Salvaterra, 1999). Majority of AL local neurons are GABAergic which receives projections both from OSN as well as PN and are known to modulate PN output via GABA receptors (Wilson and Laurent, 2005). GABAergic connections of LN onto the dendrites of PNs in AL are known to synchronise the PNs output and regulate the changes in MB neurons further upstream in higher brain centres (Laurent et al., 1998). Habituation might be the result of either depression of excitatory neurotransmission (Glanzman, 2009) or potentiation of inhibitory transmission (Das et al., 2011; Paranjpe et al., 2012). We asked how ChT contributes to the devaluation of inconsequential stimuli, through excitatory cholinergic or inhibitory GABAergic. In such a scenario, there could be two possible mechanisms: 1) ChT might be present in GABAergic neurons and directly regulate olfactory habituation, 2) ChT might regulate ACh release from excitatory cholinergic neurons which in turn activate GABAergic neurons for inhibition. Therefore, we assessed both the possible mechanisms and assessed if ChT is required in GABAergic or cholinergic neurons to facilitate olfactory habituation learning.

The synthesis of the GABA neurotransmitter is catalysed by enzyme Glutamic acid decarboxylase (GAD). *GAD1GAL4* has a predominant expression in inhibitory GABAergic neurons in the fly brain (Okada et al., 2009). We used *GAD1GAL4* to mark the putative GABAergic neurons in the larval brain by driving *UAS-mCD8GFP* and assessed if ChT is colocalised in the GABA positive regions. A superposition of GABA positive neurons and ChT was observed in MB calyx and subesophageal ganglia (SEG) (Fig. 5 A-I). This was an interesting observation because ChT is a transporter generic to ACh metabolic cycle, but we observed a major ChT immunopositive region colocalised with GABAergic terminals in larval central brain neuropile. To elucidate whether ChT is indeed required in inhibitory neurons for olfactory habituation, we knockdown ChT by expressing *UAS-ChT^RNAi1^* and *UAS-ChT^RNAi2^* in GABAergic neurons using *GAD1*GAL4. Knockdown of ChT by both *ChT^RNAi1^* and *ChT^RNAi2^* significantly reduced chemotaxis as well as habituation (Fig. 5 J, K). The waning of the response towards odour stimulus was recovered back in the control group of larvae (*GAD1*GAL4>+) after 30 min rest time or by giving cold stimulus but not in ChT depleted group of larvae (*GAD1GAL4>UAS-ChT^RNAi1^* and *UAS-ChT^RNAi2^*) (Fig. 5 L). The habituation ability was restored to baseline levels on transgenic expression of ChT in a *ChT^RNAi1^* background in genotype GAD*1GAL4>ChT^RNAi1^;UAS-ChT* (Fig. 5L). These observations suggest that ChT has a function in GABAergic neurons which is necessary for olfactory habituation. To further ascertain that the habituation defect observed due to knockdown of ChT in GABAergic neurons is an acute effect and not developmentally driven defect, we genetically restricted the knockdown of ChT in GABAergic neurons to the shorter developmental window of 3^rd^ instar foraging larval stage using *TubGAL80^ts^*. Significantly enhanced chemotaxis was observed when knockdown of ChT was restricted to the specific developmental window of 3^rd^ instar foraging larvae at 29°C but not at 18°C (Fig 5M). However, when we knocked down ChT under *TubGAL80^ts^* throughout development, we observed the reduction in naïve chemotaxis (Fig 5M) which was similar to our previous observation with *GAD1GAL4>ChT^RNAi1^* and *GAD1GAL4>ChT^RNAi2^* group of larvae (Fig. 5J). The habituation index was also significantly reduced when ChT knockdown was specified to the developmental window of 3^rd^ instar foraging larvae using *TubGAL80^ts^* at 29°C but not at 18°C (Fig. 5N). However, the reduction was more drastic when it was knocked down throughout development (Fig. 5N). These experiments allowed us to differentiate the acute effect of ChT knockdown upon chemotaxis and habituation from the developmental role of these neurons in the expression domain of GAD1. Together, these results show that ChT is an essential component of inhibitory GABAergic neurons and facilitate olfactory habituation directly.

**Figure 5:**
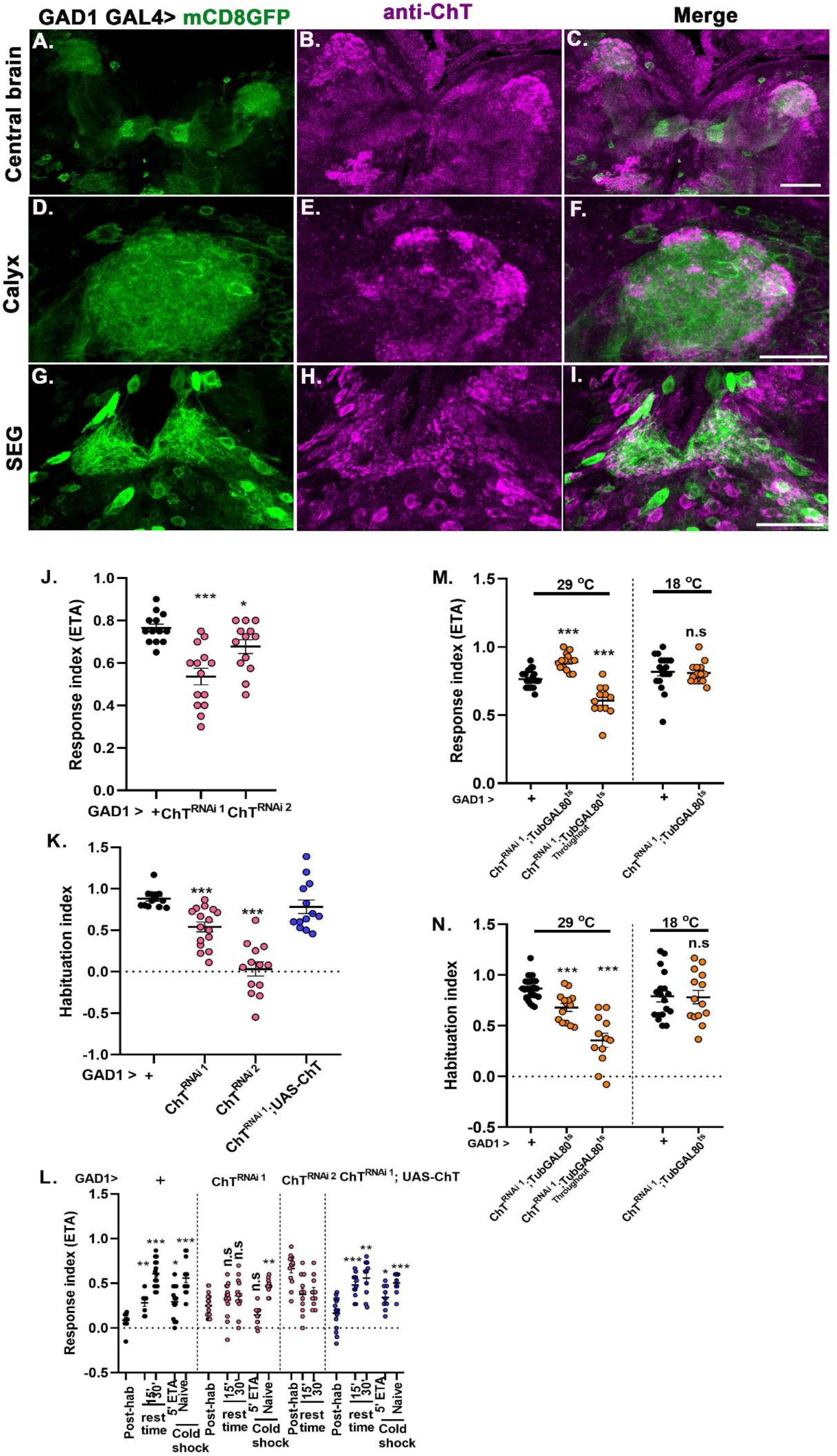
ChT is localised to GABAergic neurons and has a functional role in habituation as well as chemotaxis. (A-I) shows mCD8GFP driven by *GAD1GAL4* (green), coimmunostained with anti-ChT (magenta) and merge (colocalization appears as white). Staining marks several areas of central brain (A-C) including calyx (D-F) and subesophageal ganglia (SEG, G-I). All immunostained images are representative of 3-5 brains and scale bar is 50 μm. (J and K) shows response index and habituation index, of larvae with genotype *GAD1GAL4*> *+* (black circles, N≥13), *GAD1GAL4> ChT^RNAi1^ and ChT^RNAi2^ (pink circles,* N≥13), *GAD1*GAL4> *ChT^RNAi1^; UAS-ChT* (blue circles, N≥17). (L) Response indices towards ETA of larvae after exposure to 5 min ETA (post-hab), in absence of stimulus after 15 and 30 min rest-time, larvae presented to 1 min cold shock (5’ETA+coldshock), naïve larvae presented to 1 min cold shock (naïve + cold shock) in genotypes *GAD1GAL4*> *+* (black, N≥13), *GAD1GAL4> ChT^RNAi1^ and ChT^RNAi2^* (pink, N≥12)*, GAD1GAL4*> *ChT^RNAi1^;UAS-ChT (Blue,* N≥15). (M) response index, (N) habituation index of larvae towards ETA by ChT knock-down at the specific developmental window of 3^rd^ instar foraging larvae using temperature controlled GAL80^ts^ in genotypes *GAD1GAL4*> *ChT^RNAi1^;TubGAL80^ts^* (orange circles) at 29 °C (N=14), throughout 29 °C (orange circles, N=12) and at 18 °C (orange and black circles, N≥14) as compared to *GAD1GAL4*>+ (Black circles, N=24). Statistical significance determined using Mann-Whitney U test, *** represent p≤0.0001, n.s represent p≥0.05

Next, we knocked down ChT in cholinergic excitatory neurons which also includes excitatory interneurons and PNs of the olfactory pathway using *ChATGAL4* (Salvaterra and Kitamoto, 2001). Null mutant alleles of ChAT encoding genes in *Drosophila* leads to embryonic lethality and show morphological abnormalities suggesting that ACh mediated neurotransmission is important for development (Kitamoto et al., 2000). Therefore, to bypass any developmental defect, we knocked down ChT in the specific foraging 3^rd^ instar larval developmental window. A drastic reduction of chemotaxis was observed when ChT was knocked down in genotypes *ChATGAL4>ChT^RNAi1^ or ChT^RNAi2^* (Fig. 6 A). However, enhanced chemotaxis or hypersensitivity was observed when ChT knockdown was restricted to foraging 3^rd^ instar larvae stage in genotype *ChATGAL4>ChT^RNAi1^;TubGAL80^ts^* at 29°C (Fig. 6 B). Interestingly, we saw habituation defect only when ChT was knocked down throughout the development in genotype *ChATGAL4>ChT^RNAi1^;TubGAL80^ts^* at 29°C but not when specifically knocked down during foraging 3^rd^ instar larvae (Fig. 6C). The reduced chemotaxis, as well as habituation upon knockdown of ChT throughout development, might have arisen due to modulation of olfactory neural circuit development which is dependent on the ACh mediated signals.

**Figure 6:**
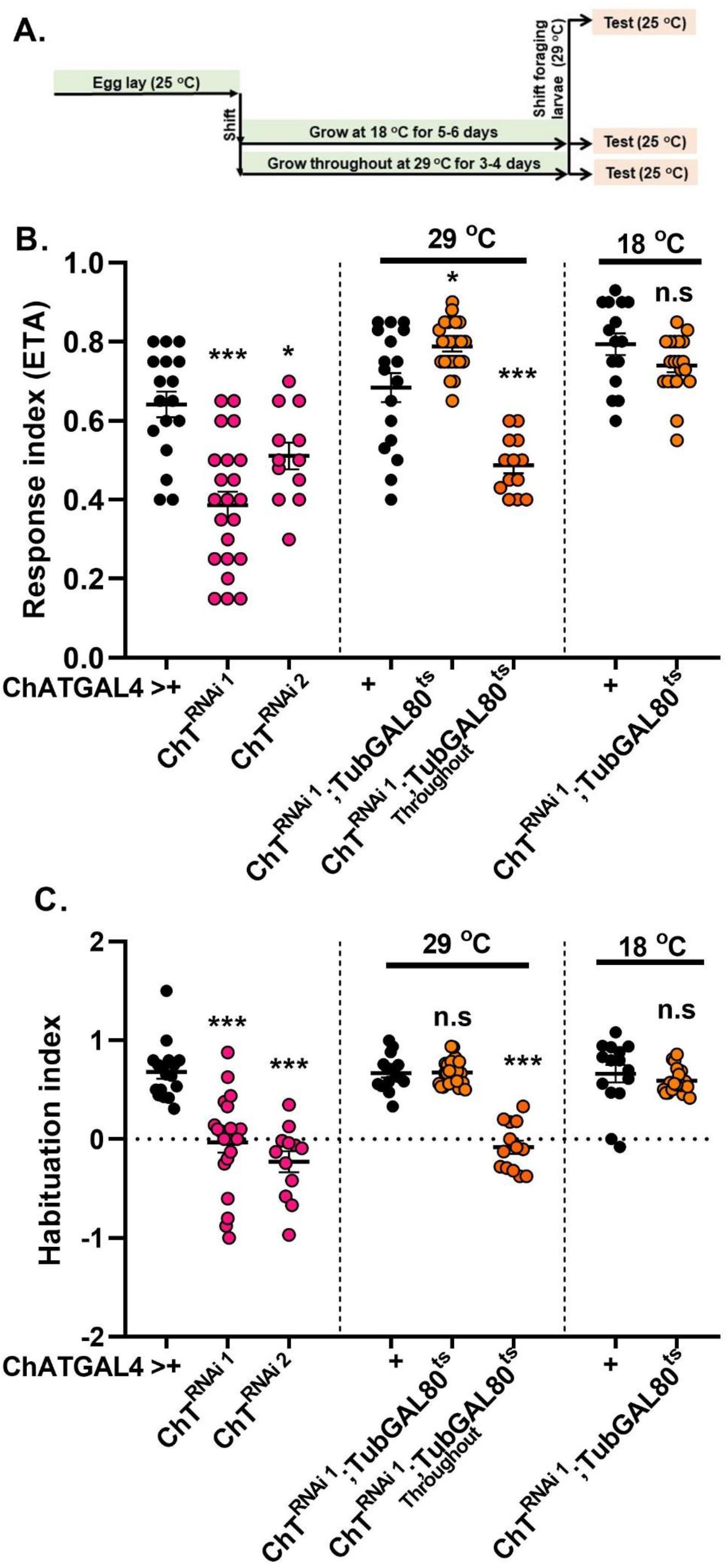
Knockdown of ChT in cholinergic neurons in specific developmental window does not alter habituation but show enhanced chemotaxis. (A) Schematics represent the specific time segments followed for the experiments. (B) response index, (C) habituation index of foraging larvae towards ETA in genotypes *ChATGAL4*>+ (black circles, N=13), *ChAT*GAL4> *ChT^RNAi1^ or* ChT^*RNAi2*^ (pink circles, N≥13), *ChATGAL4>ChT^RNAi1^;TubGAL80^ts^* at 29°C where ChT was knocked down only in 3^rd^ instar foraging developmental window, throughout 29°C (orange circles, N≥13) and at 18°C (orange and black circles, N≥13) as compared to *ChATGAL4*>+ (black circles, N≥13). Statistical significance using Mann-Whitney U test. *** represent p≤0.0001, n.s represent p≥0.05.

**Figure 7:**
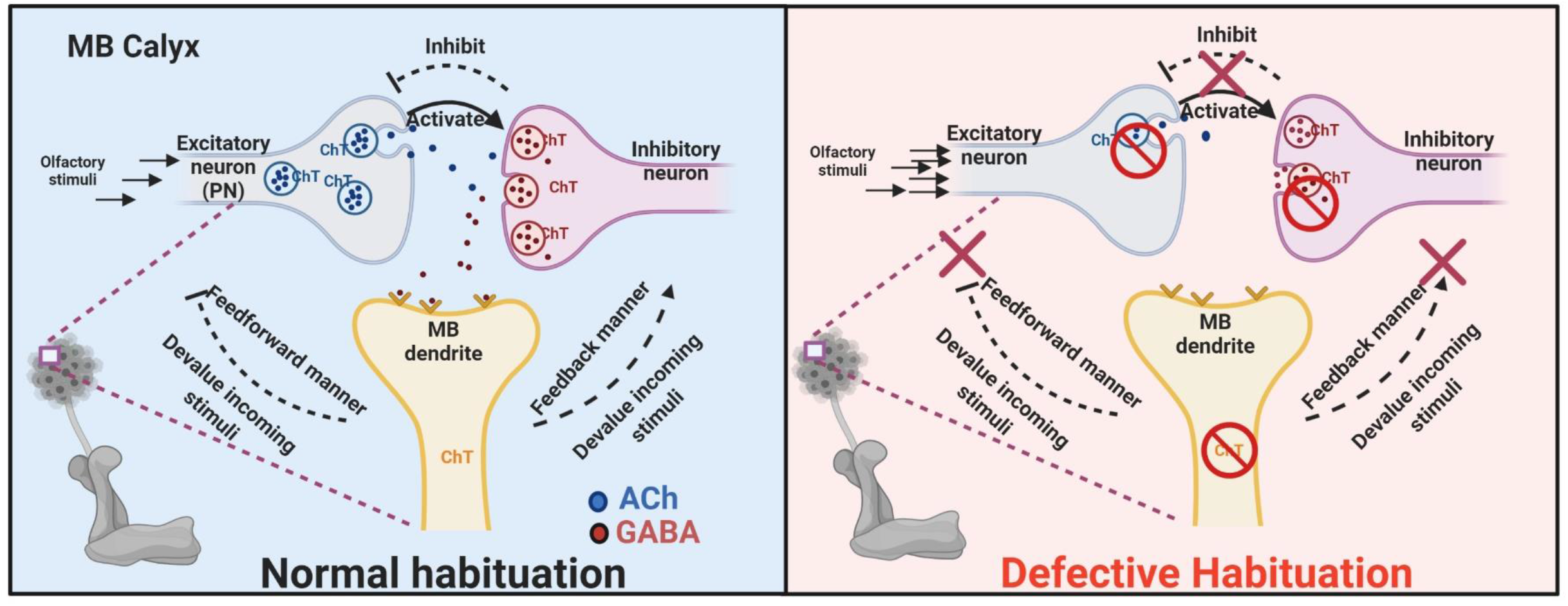
A functional model of neuronal subsets localising ChT at glomerulus of MB calyx to regulate olfactory habituation. *Left panel*, PN carries olfactory information to the calyx of MB and activate inhibition of MB neurons via activation of GABAergic inhibitory neurons. Both excitatory PN and inhibitory GABAergic neurons requires ChT for the forward inhibition and permits control over the MB neurons to respond to input stimuli. There are two possibilities through which ChT in MB may further manifest control over incoming stimuli as depicted in the figure (black dashed lines) : a feedback inhibition of incoming stimuli via GABAergic neurons which inturn may inhibit output from excitatory PN or a feedforward inhibition of incoming stimuli. *Right panel,* if ChT in PN, GABAergic or MB intrinsic neuron is depleted (shown as red no-sign icon), the habituation is defective and the control over the incoming stimuli is lost leading to enhanced chemotaxis towards odour, as presented in our results.

Together, our results suggest that ChT mediates regulation of chemotaxis by a subset of both inhibitory and excitatory cholinergic neurons but facilitate habituation only through the inhibitory GABAergic neurons. These observations strongly suggest that ChT has a function both in inhibitory as well as excitatory neurons. The attenuated chemotaxis and habituation upon selective knockdown of ChT in GABAergic neurons indicate that ChT might not be unique to cholinergic neurons but may facilitate the functioning of other neurotransmitter system too.

## Discussion

In this study, we demonstrate ChT dependent plasticity in the olfactory circuit and higher processing centres to devalue the incoming olfactory stimuli (habituation), followed by spontaneous recovery in absence of the stimuli as well as recovery upon exposure to unrelated stimuli (dishabituation) from a habituated state. We show that hypersensitivity is a direct corollary of habituation defects due to depletion of ChT in MB. ChT function has classically been attributed to acetylcholine formation. Importantly, we report for the first time that ChT is also present in GABAergic neurons and contribute to habituation and sensory stimuli suppression. Thus, in addition to highlighting the importance of ChT in attaining habituation flexibility, this data establishes a relationship between upstream plasticity and downstream stimulus which is mediated by ChT localised in MB and GABAergic neurons.

The requirement of bout control over perceived sensory stimuli is important for an animal to acquire transition flexibility between a habituated to non-habituated state and back which enables an animal to respond selectively to a stimulus depending on the demand. Our results show that ChT is one, if not an exclusive, regulatory molecule to facilitate this flexibility. This is supported by our observations that ChT is required for habituation, spontaneous recovery, dishabituation and stimulus suppression.

We propose that ChT functions as a regulatory switch in MB to control the perceived olfactory signals in order to facilitate habituation behaviour. This impetus is lost when ChT is depleted in MB leading to hypersensitivity. MB lesion or ablation in *Drosophila* show elevated locomotor activity (Heisenberg et al., 1985) and diurnal activity (Martin et al., 1998). Recently conditional silencing of MB output neurons (MBON) was reported to show enhanced proboscis extension reflex (Chia and Scott, 2020). Knockdown of dopaminergic receptors (DAMB) in MBON-1 ped>αβ also promotes yeast food-seeking behaviour in fed flies as well as odour seeking behaviour (Tsao et al., 2018). Our finding that lack of ChT in MB causes hypersensitivity corroborates with previous behavioural responses on ablation of MB in diverse insects suggesting ChT may contribute to such elicited phenotypes. Although our results do not provide any direct proof and require further investigation in an anatomical context of how ChT in MB controls incoming stimuli, it might likely be occurring via feedback regulation from MB toMB calyx or via feedforward from MB to AL. Several anatomical evidence support feedback regulation in sensory processing via connections from AL to OSN (Olsen and Wilson, 2008), MB to PN (Tanaka et al., 2009), MB to MB calyx in *Drosophila* (Masuda-Nakagawa et al., 2014), cortex to the olfactory bulb in mammals (Balu et al., 2007), cortex to visual centres in mammals (Tiesinga et al., 2008) or feedforward regulation from MB to AL in *Drosophila* and honeybees (Hu et al., 2010; Kirschner et al., 2006).

Genetic experiments demonstrate olfactory habituation in *Drosophila* as GABA mediated inhibition of PN terminals at AL which required NMDA receptors or GABA receptors on dendrites of cholinergic PN in AL (Das et al., 2011; Paranjpe et al., 2012). Intriguingly, we observed that ChT is localised at both cholinergic and GABAergic terminals of MB calyx and functionally required for olfactory habituation. Immunoelectron microscopic analysis of MB calyx reveals that each glomerulus comprises cholinergic nerve endings at the core which is encircled by a numerous GABAergic terminal (Yasuyama et al., 2002). Therefore, the dendrites of MB Kenyon cells receive major excitatory inputs via cholinergic and inhibitory from GABAergic elements which exist in close vicinity with each other. Such anatomy has been reported in many invertebrates such as honeybees (Ganeshina and Menzel, 2001), crickets (Schurmann et al., 2008), locusts (Leitch and Laurent, 1996) in addition to *Drosophila*. The localisation of ChT in AL and MB calyx glomerulus and the fact that both the locations have excitatory as well as inhibitory terminals in close vicinity indicate that ChT might be important for the functioning of local transmission of neurotransmitters. We speculate that ChT is a conserved protein and its presence in both excitatory and inhibitory nerve terminals in the glomerulus of MB calyx and AL might represent a conserved phenomenon across species. Certainly, more specific studies are required of ChT function in different neurotransmitter systems to understand the functional diversity of this transporter. Based on our observations, we propose a mechanistic model that represents how ChT might mediate habituation and stimulus suppression by a synchronised activity between excitatory, inhibitory and MB neurons and maintains a balance between perceived stimuli and habituation plasticity (Fig. 7).

Multiple diverse phenotypes are attributed to ASD (Sinha et al., 2014). It is unclear if all of these phenotypes share any single causative factor. Unusual sensory processing experienced as overwhelming or augmented sensory perceptions across different sensory modalities is a common phenotypic trait of ASD (Bonnel et al., 2003; Leekam et al., 2007; Tavassoli et al., 2014). This hypersensitivity has been described to be closely associated with lack of habituating capability towards a sensory stimulus. Previous evidence suggests that attenuated habituation is quite prevalent in people with ASD and may play a role in unusual sensory sensitivity (Guiraud et al., 2011; Kleinhans et al., 2009; Webb et al., 2010). Besides, ASD animal model studies and human observations also suggest dysfunction of GABA signalling (Cellot and Cherubini, 2014). Also, ChT polymorphism associated with attention deficit hyperactivity disorder (ADHD) has been reported as a key trait of Autism (English et al., 2009). Integrating all the phenotypes (i.e defective habituation, hypersensitivity, the role of inhibitory signalling) obtained in this study by knockdown of ChT in MB and GABAergic neurons projects to a unifying view of functional relatedness between ChT and multiple salient phenotypes of ASD. Our results provide a cohesive perspective of ChT as one of the common cause underlying multiple aspects of Autism and provide avenues for future research to determine neural or molecular correlates of ChT for developing improved diagnosis and therapy.

## Acknowledgements

We thank Dr Rakesh K Mishra for his constant support and guidance during the completion of the work. We thank CCMB for providing lab space, granting access to central imaging and fly facility and necessary infrastructure support. We thank Dr Gaurav Das for critical reading of the manuscript and providing insightful comments. We also thank Dr Sonal Nagarkar Jaiswal, Nikhil Hajirnis and Shagufta Khan for the scientific discussions and useful suggestions. We also acknowledge the assistance received from Dr V Bharathi and associated dissertation trainees at various stages of completion of this work. We acknowledge BDSC, VDRC, NCBS fly facility and Dr. Gaurav Das for providing fly stocks. This work was supported by the research grant of the Department of Science and Technology, Govt. of India vide reference no. SR/CSRI-P/2017/2 to Runa Hamid under Cognitive Science Research Initiative Scheme.

## Author contribution

Conceptualisation, funding acquisition, project administration, formal analysis and manuscript preparation by R.H. Data Curation by R.H and H.S.S. Standardisation and preliminary data generation by R.H and M.N.K.

## Supplementary figures

**Figure S1:**
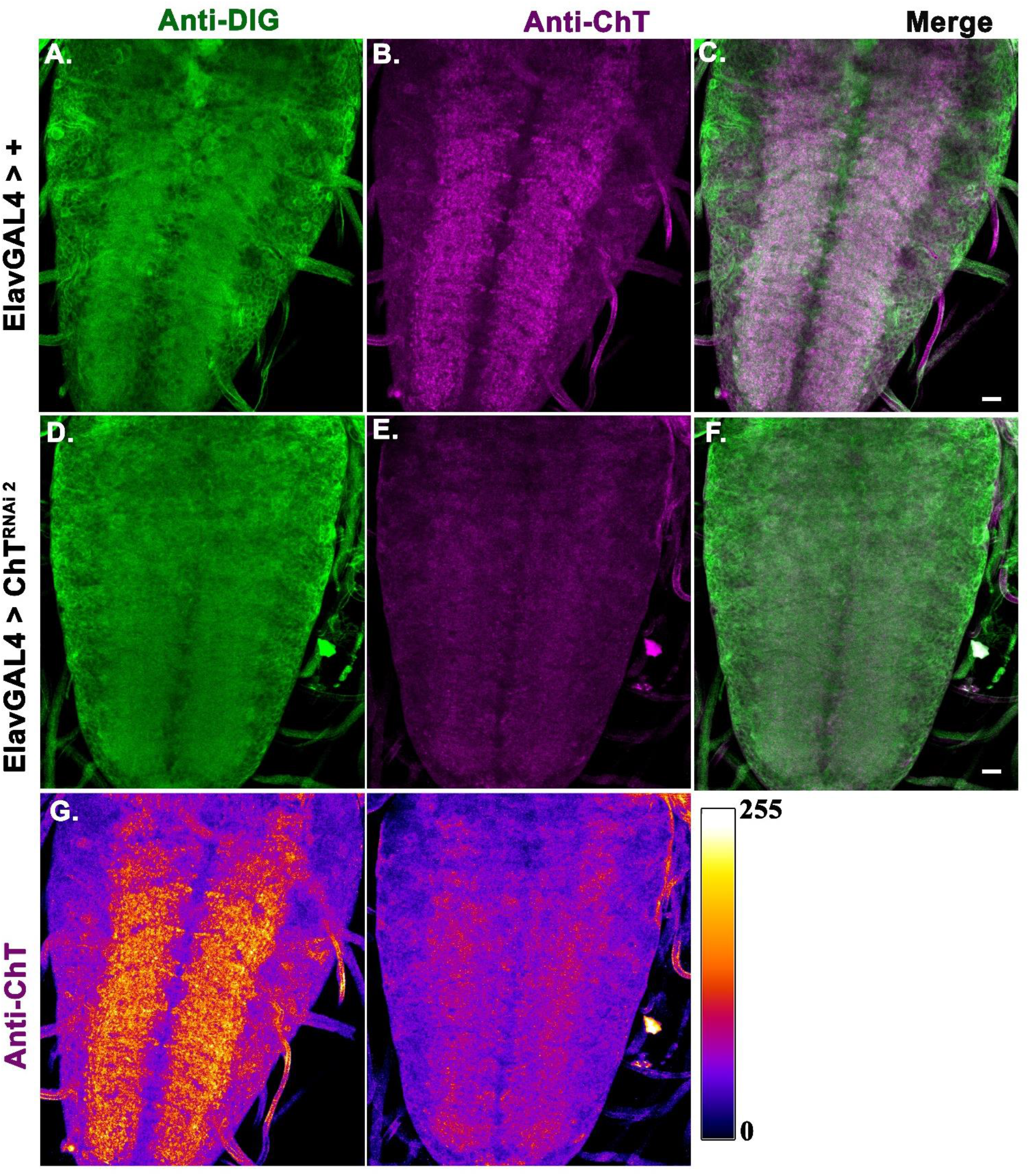
RNAi mediated knockdown of ChT using ChT^RNAi2^shows reduction in protein levels of ChT in larval VNC by immunostaining. (A-C) Immunostaining with anti-Dlg (green), anti-ChT (magenta) and colocalised region shown as white in merge image in genotype *Elav;;dicer>+* compared to (D-F) with genotype *Elav;;dicer>UAS-ChT^RNAi2^*. (G) Images of VNC immunostained with *anti-ChT* of *Elav;;dicer>+ (left)* and *Elav;;dicer>ChT^RNAi2^ (Right)* converted to Fire LUT map using imageJ. The scale shows the range of colours from 0-255pixel intensity. These are representative images of 3-5 brains. Scale bar 50μm.

**Figure S2:**
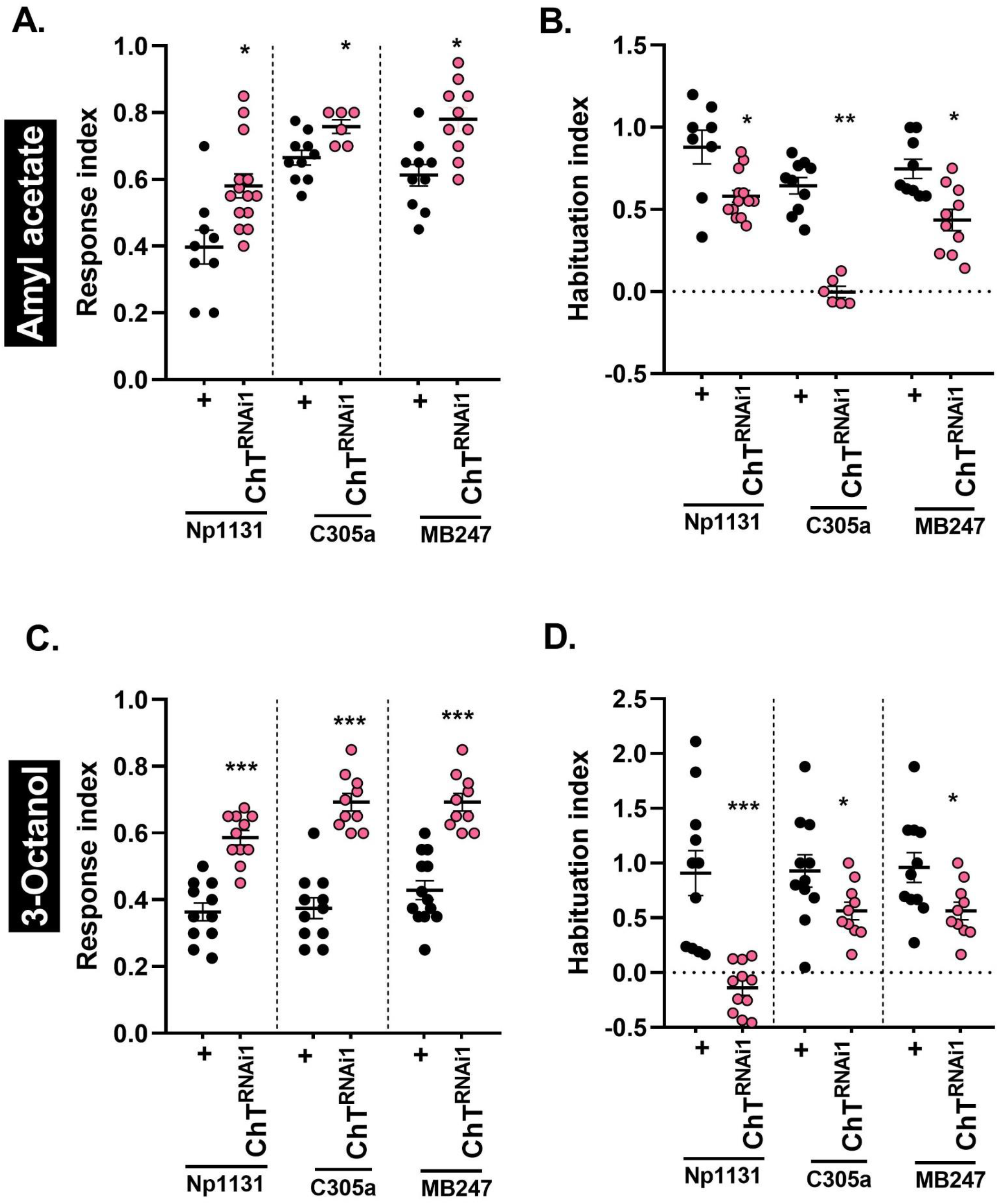
Knockdown of ChT in MB intrinsic neurons attenuates chemotaxis and habituation irrespective of the kind of odour: (A and B) Response index and habituation index (B) towards amylacetate (AMA, 1:100 dilution) of genotypes *NP1131>ChT^RNAi1^* (pink circles, N=14) as compared to *NP1131> +* (black circles, N=9), *C305aGAL4>ChT^RNAi1^* (pink circles, N=6) as compared to *C305aGAL4> + (*black circles, N=10) *and MB247GAL4>ChT^RNAi1^* (Pink circles, N=10) as compared to *MB247>+ (*black circles, N=10). (B and D) Response index and habituation index towards 3-Octanol (3-Oct, 1:1000 dilution) of genotypes *NP1131>ChT^RNAi1^* (Pink, N=11) as compared to *NP1131> +* (Black, N=11), *C305aGAL4>ChT^RNAi1^* (Pink, N=10) as compared to *C305aGAL4> +* (Black, N=9) *and MB247GAL4>ChT^RNAi1^* (Pink, N=10) as compared to *MB247>+* (Black, N=13). Data shown as scatter plot and statistical significance was determined by Mann-Whitney U test. ***p<0.0001

**Figure S3:**
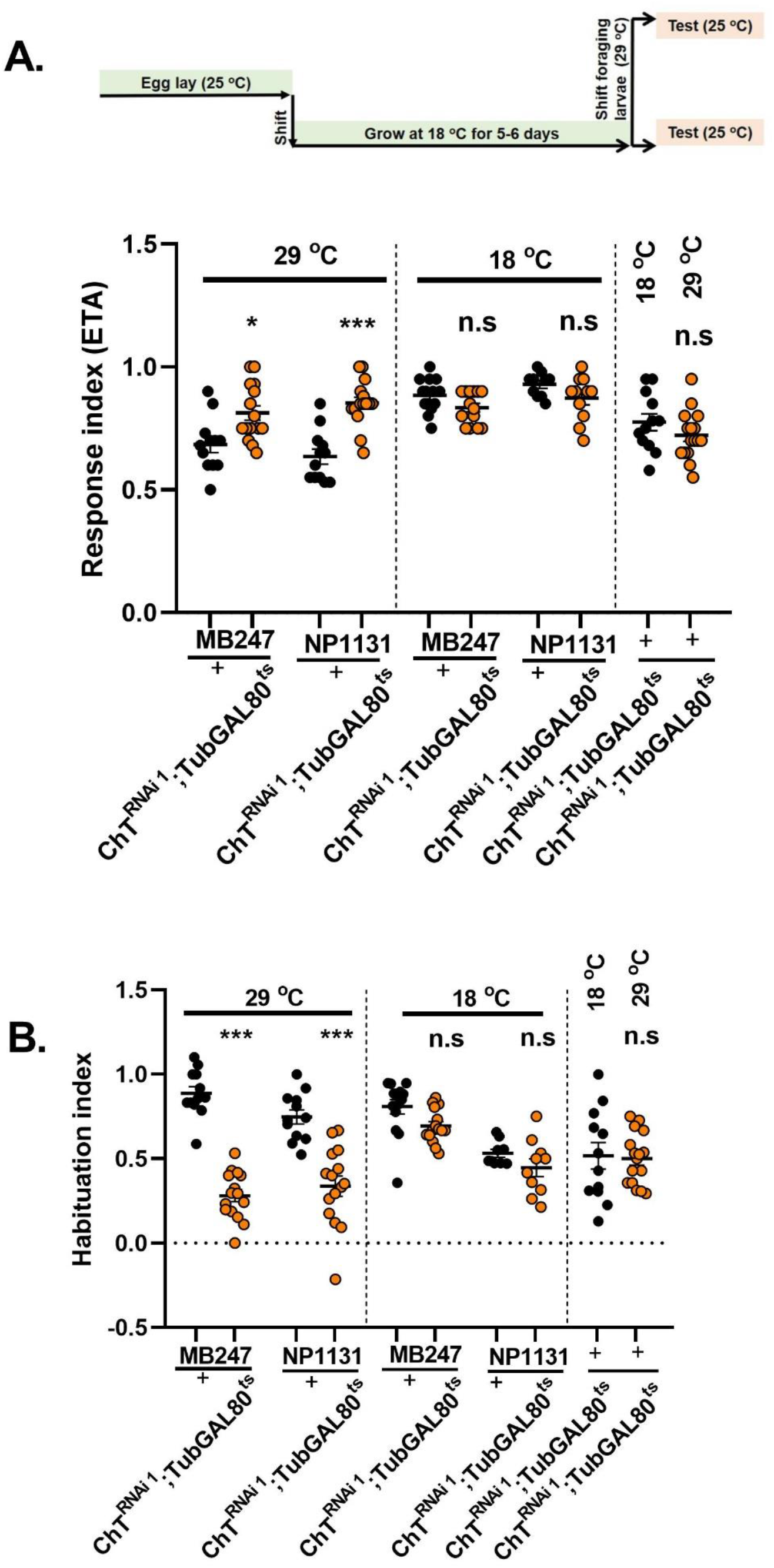
Enhanced chemotaxis and reduced habituation is an acute effect of ChT knock down. A) Schematic shows specific time window of temperature shift for TARGET system using *TubGAL80^ts^*. Response index of group of larvae with genotype *MB247GAL4*>*ChT^RNAi1^;TubGAL80^ts^* (orange circles) compared to control *MB247GAL4*>*+* (Black circles, N≥12), *NP1131GAL4*>*ChT^RNAi1^;TubGAL80^ts^* (orange circles) compared to control *NP1131GAL4*>*+* (Black circles, N≥12) at 29°C and 18°C, *ChT^RNAi1^;TubGAL80^ts^* > + at indicated temperatures of 29°C and 18°C (N≥12). B) Habituation index of larvae of *MB247GAL4*>*ChT^RNAi1^;TubGAL80^ts^* (orange circles) compared to control *MB247GAL4*>*+* (Black circles, N≥12), *Np1131GAL4*>^*ChTRNAi1*^;TubGAL80^*ts*^ (orange circles) compared to control *NP1131GAL4*>*+* (Black circles, N≥12) at 29°C and 18°C, and *ChT^RNAi1^;TubGAL80^ts^* > + at indicated temperatures of 29°C and 18°C (N≥12). Statistical significance using Mann-Whitney U test*** represent p≤0.0001, n.s represent p≥0.05.

**Figure S4:**
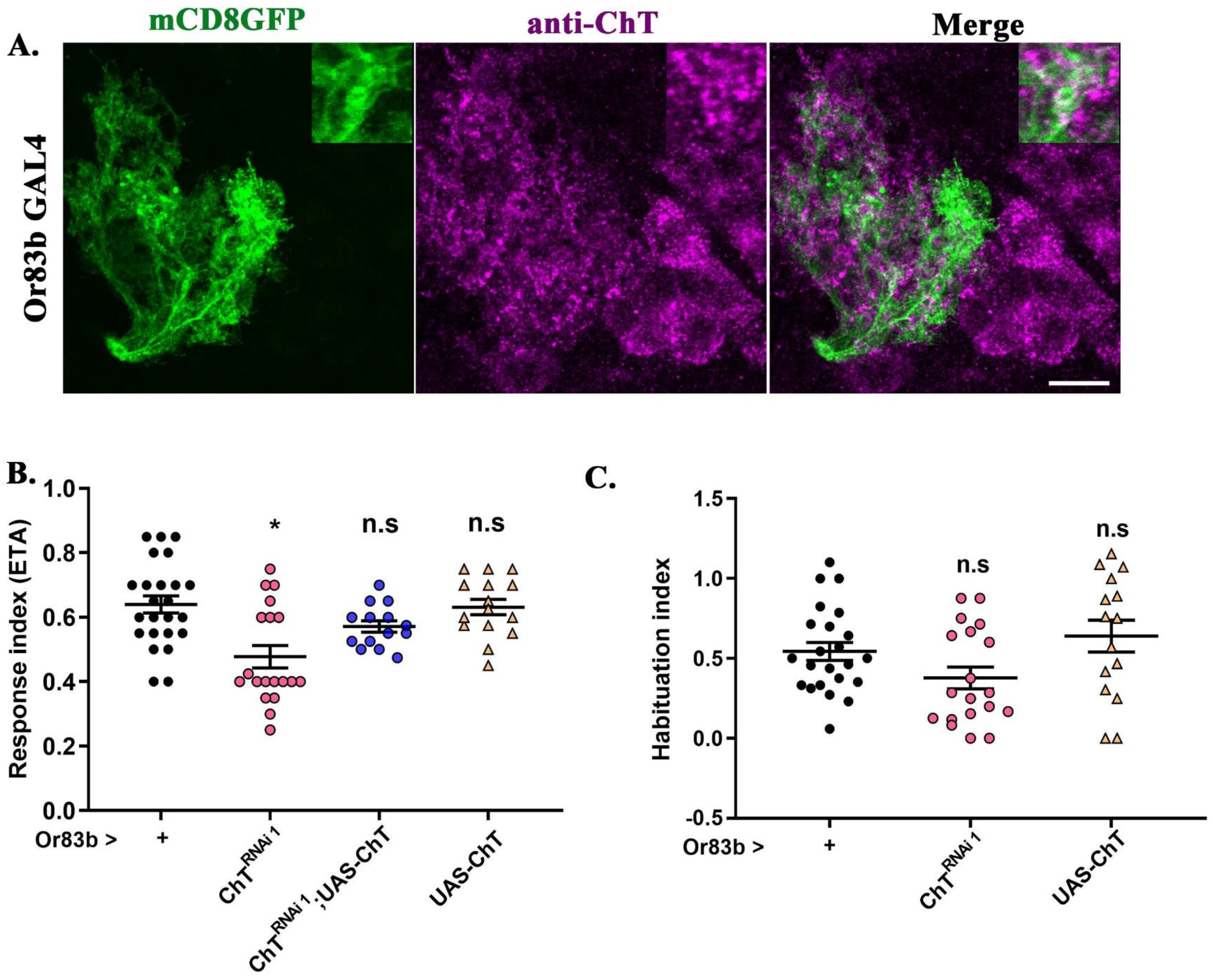
Knockdown of ChT in olfactory sensory neurons does not affect habituation but attenuates chemotaxis. (A) Immunostained images of larval olfactory sensory neurons in 3^rd^ instar dissected larval brain marked by expression of mCD8GFP using *Or83bGAl4* driver in genotype *Or83bGAL4>UAS-mCD8GFP* (green), costained with anti-ChT (magenta), merge (colocalised regions of mCD8GFP and ChT appear as white). Inset shows a cropped and zoomed image of colocalised (mCD8GFP and ChT) terminal of OSN. Image shown is a representative image of 3-5 brain lobes. Scale bar 50 μm (B) Response index, and (C) habituation index towards ETA of genotypes *Or83bGAL4>+* (black circles, N=23), *Or83bGAL4>ChT^RNAi1^* (pink circles, N=19), *Or83bGAL4>ChT^RNAi1^;UAS-ChT* (blue circles, N=18) and *Or83bGAL4>UAS-ChT* (yellow triangles, N=14). Statistical significance was determined by Mann-Whitney U test. * p≤0.01, n.s is non-significant where p≥0.05

**Figure S5:**
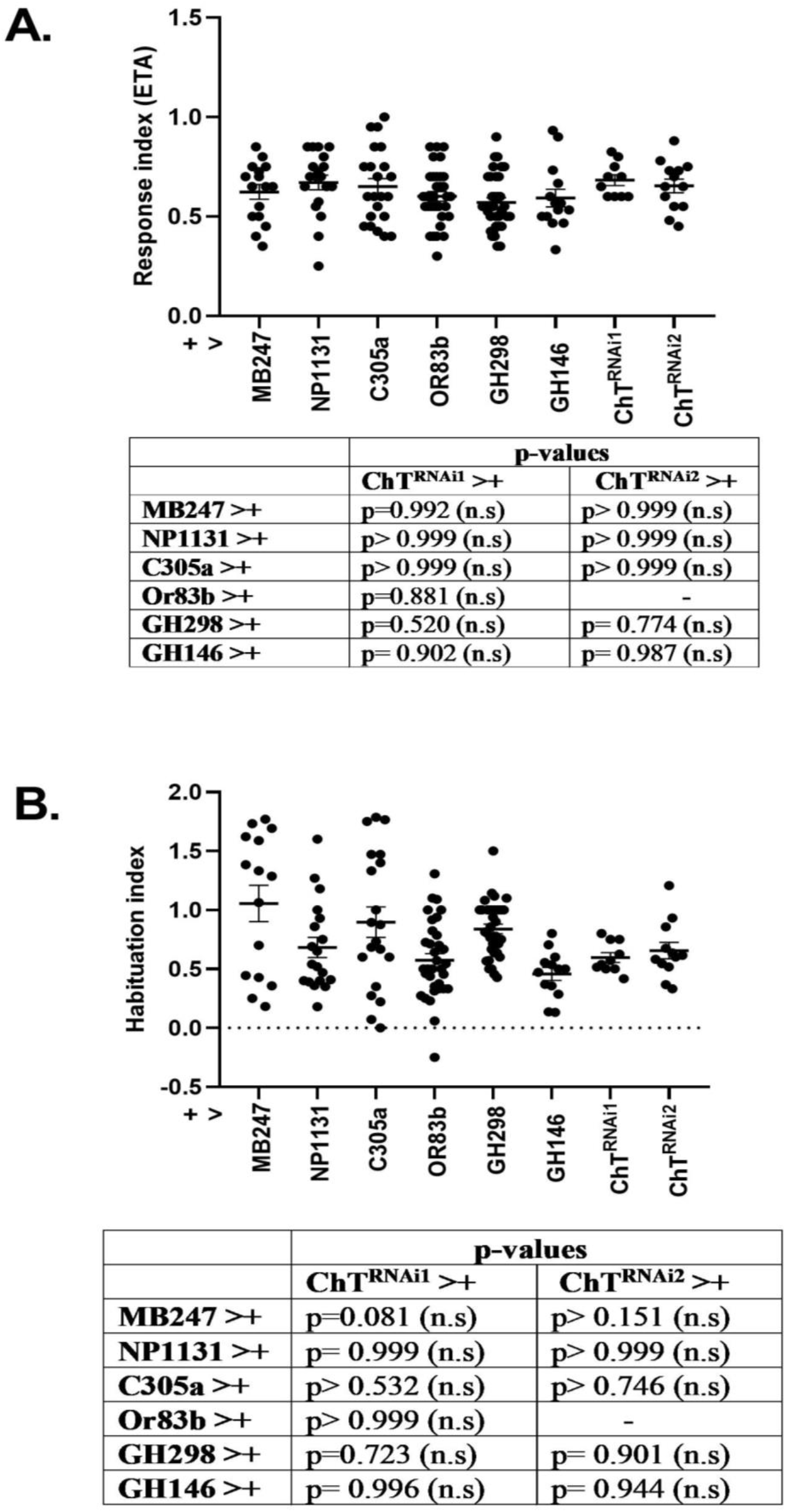
Response index and habituation index of larvae in heterozygous combinations of different *GAL4* lines versus *ChT RNAi* lines. Scatter plot showing (A) Response index (B) habituation index, of larvae with genotypes *MB247GAL4>+, NP1131GAL4>+, C305aGAL4>+, Or83bGAL4>+, GH298GAL4>+, GH146GAL4>+* as compared with *UAS-ChT^RNAi1^>+* and *UAS-ChT^RNAi2^*. Tables show respective p-values for each comparison. Statistical significance was determined using one-way Analysis of variance. n.s represent non-significance.

